# Live cell imaging reveals that RecA finds homologous DNA by reduced dimensionality search

**DOI:** 10.1101/2020.02.13.946996

**Authors:** Jakub Wiktor, Arvid H Gynnå, Prune Leroy, Jimmy Larsson, Giovanna Coceano, Ilaria Testa, Johan Elf

**Author notes:** equal contribution (Jakub Wiktor, Arvid H Gynnå). **Materials & Correspondence**. All strains, raw data and analysis codes will be provided upon reasonable request to J. E.

## Abstract

Homologous recombination (HR) is essential for the accurate repair of double-stranded DNA breaks (DSBs); it begins when the RecBCD2 complex resects the ends of the DSB into 3' single-stranded DNA (ssDNA) on which a RecA filament assembles. HR depends on the ability of this RecA-ssDNA filament to locate the homologous repair template on the sister chromosome. The mechanism by which the homology is located among vast amounts of heterologous DNA is not yet understood, despite a long history of research. Here, we directly visualize the repair of DSBs in hundreds of individual cells, using high-throughput microfluidics and fluorescence microscopy. We find that in E. coli, DSB repair is completed in 15 minutes without fitness loss. We further show that the search takes less than 10 minutes and is mediated by a thin, highly dynamic RecA filament that stretches throughout the cell. We propose a model in which the architecture of the RecA filament effectively reduces search dimensionality to two dimensions. The model is corroborated by the observation that the search time does not depend on the length of the cell or the amount of DNA, and also predicts a search time that is consistent with our measurement. Since the RecA family proteins are conserved in all organisms, our results also translate to other systems that rely on homologous recombination.

## Introduction

Homologous recombination (HR) is essential for the accurate repair of double-stranded DNA breaks (DSBs)^1^; it begins when the RecBCD^2^ complex resects the ends of the DSB into 3’ single-stranded DNA (ssDNA) on which a RecA filament assembles^3^. HR depends on the ability of this RecA-ssDNA filament to locate the homologous repair template on the sister chromosome^4^. The mechanism by which the homology is located among vast amounts of heterologous DNA is not yet understood, despite a long history of research. Here, we directly visualize the repair of DSBs in hundreds of individual cells, using high-throughput microfluidics and fluorescence microscopy. We find that in *E. coli*, DSB repair is completed in 15 minutes without fitness loss. We further show that the search takes less than 10 minutes and is mediated by a thin, highly dynamic RecA filament that stretches throughout the cell. We propose a model in which the architecture of the RecA filament effectively reduces search dimensionality to two dimensions. The model is corroborated by the observation that the search time does not depend on the length of the cell or the amount of DNA, and also predicts a search time that is consistent with our measurement. Since the RecA family proteins are conserved in all organisms^5^, our results also translate to other systems that rely on homologous recombination.

To study the mechanism of the homologous recombination directly in living bacteria, we created an inducible DSB reporter system consisting of (**i**) an inducible Cas9 nuclease to create DSBs at a single, specific chromosomal location (referred to as ‘*cut-site*’), (**ii**) a fluorescent *parS*/mCherry-ParB system (referred to as ‘ParB’) to visualize the chromosomal location of the break using fluorescence microscopy, and (**iii**) an SOS-response^6^ reporter to select the cells undergoing DSB repair (**Fig. 1a, Fig. S1b**). We used a variant of the microfluidic mother-machine device^7^ that allows for fast, accurate, and automated induction of Cas9 nuclease (**Fig. 1a, Fig. S1b**).

**Figure 1.**
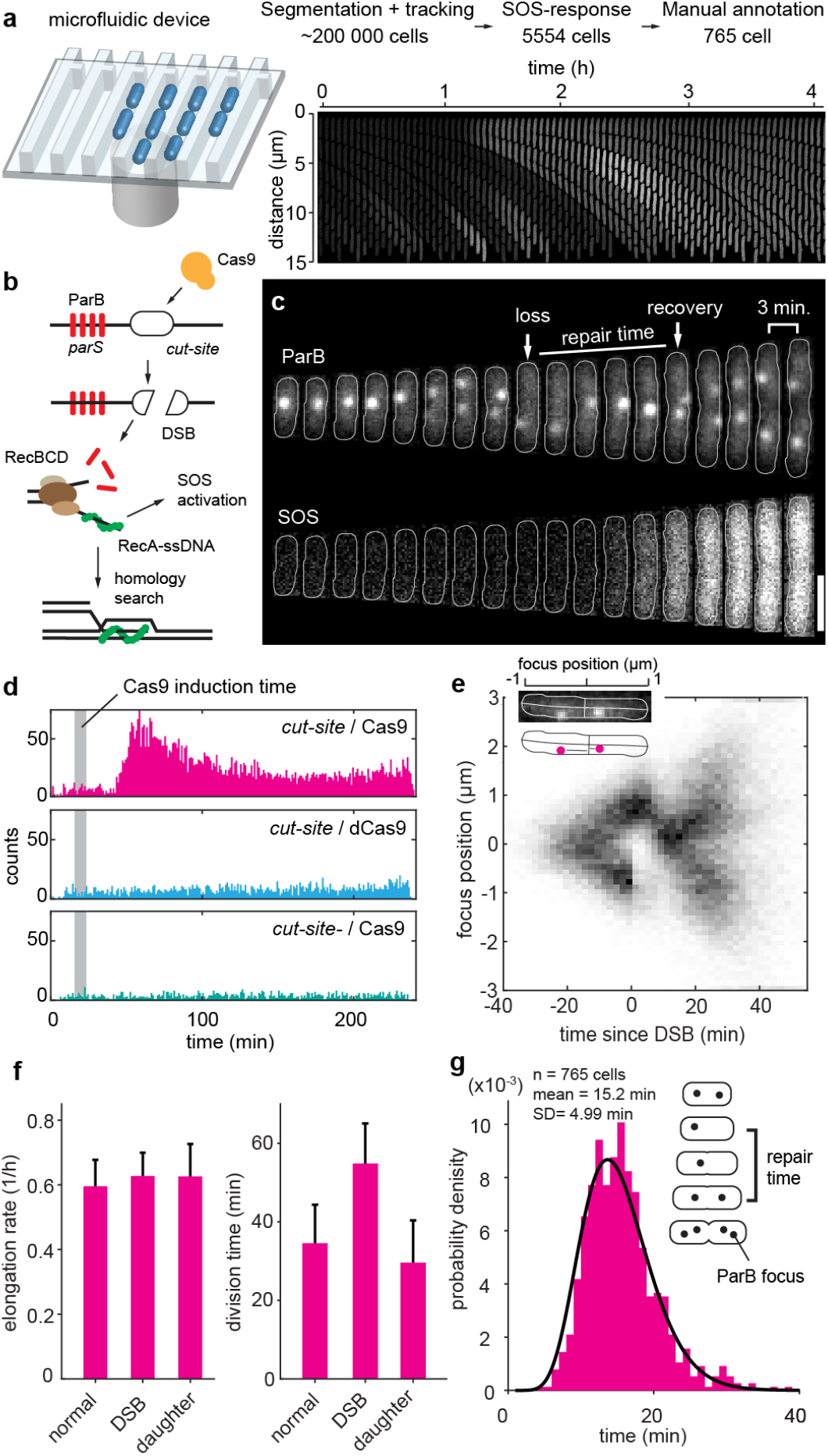
High throughput imaging of DSB repair. a) *Left*: DSB induction experiments. Cells were grown in a mother machine microfluidic device. Cells were segmented, tracked, selected based on the increase in fluorescence from a plasmid SOS reporter, and finally the ParB loss and recovery were manually annotated. The number of cells processed in each step is shown. *Right*: montage of a single growth channel showing SOS activation after induction of DSBs. b) Cartoon showing formation, processing, and repair of a DSB. Cas9 binds to the *cut-site* and creates a DSB. Next, RecBCD binds to DNA ends at the DSB site and begins end processing, which ejects ParB proteins. RecA is loaded on the ssDNA ends created by RecBCD. RecA bound to ssDNA induces SOS response. RecA-ssDNA filament undergoes search for homology, and after the homology is located the DSB is repaired. c) A cell undergoing DSB repair. Events of ParB focus loss and recovery are annotated, as well as the repair time. Cell outlines are displayed with a grey solid line. Scale bar is 2 µm. d) Time of half-maximum SOS reporter signal in single cells which activated SOS response, (i.e. increased CFP signal > 4-fold). Histograms for cells with *cut-site* and active Cas9 variant (5553 out of 98394 cells activated SOS-response), cut-site and dCas9 (1901 out of 98383 cells activated SOS-response), and without *cut-site* and with active variant of Cas9 (875 out of 40400 cells activated SOS-response). The grey area shows the time of Cas9 induction. e) Localization of ParB foci along the cell length during DSB repair. Cells were oriented so that the remaining ParB focus was positioned on top of the y-axis. For each cell the time was aligned based on the time of ParB focus loss, as annotated in c). Insert shows a ParB channel of a single cell overlaid with outline and backbone (top), and mapping of the foci position along the cell’s backbone (bottom). f) Average elongation rate (*left*) and division time (*right*) of normal cells, cells with a single DSB, and the daughter cells of the DSB cells. Mean and standard deviation are shown. g) DSB repair times. Solid line shows gamma (parameters: *k*= 9.98, *θ*=1.52) fit to the data.

### DSB repair is fast and accurate

Formation of a DSB is followed by end processing by RecBCD that removes ParB markers close to the break^8^, and by activation of the SOS response reporter. Both of these events can be detected using fluorescence microscopy. A pulse of Cas9 production gave rise to specific DSBs in the cells with the chromosomal *cut-site* and an active variant of Cas9 (**Fig. 1c, Fig. S2**), which was manifested by an increase in the fraction of SOS-activated cells shortly after the induction (**Fig. 1d**). Notably, expression of Cas9 in cells without the *cut-site*, or expression of catalytically dead Cas9 (dCas9), did not induce SOS response (**Fig. 1D, Fig S2b**), showing that the DSBs generated are specific. Activation of the SOS response was dependent on homologous recombination and was absent in cells lacking *recA* or *recB* genes (**Fig. S2b**). The ParB foci lost due to DSB were recovered in the wild-type cells, but not in the cells lacking critical components of recombination: *recA, recB*, or *recG* and *ruvC* (**Fig. S2c**). These results show that DSBs induced by Cas9 were repaired by HR. The DSB repair is remarkably robust: 95.5% (447/468) of the cells that retained an uncleaved template repaired the DSB and divided afterward. The repair was impaired in cells that lacked the repair template since none of the cells that cleaved all copies of the *cut-site* divided during the 4-hour experiment (n=27).

Next, we focused on the dynamics of the *cut-site* loci during the repair. Typically, after a DSB, the uncut locus first translocated to the middle of the cell and then split into two foci that segregated soon after (**Fig. 1c**). This pattern was clearly visible when we plotted the ParB foci positions along the cell length against the time relative to the DSB event (**Fig. 1e**). Next, we measured the repair times in individual cells, i.e., the time between the loss and reappearance of the ParB focus (**Fig 1c**). We limited the analysis to cells that had two separated ParB foci before the DSB. We found that a DSB can be repaired in 15.2 ± 4.99 min (mean ± SD, n = 765, **Fig. 1g**). These results were consistent between replicates (**Fig. S2**), and importantly, also when the I-SceI enzyme was used instead of Cas9 to induce breaks (**Fig. S2**). Given that the repair time is just a fraction of the generation time (here 35 ± 10 min (mean ± SD)), we asked whether it has negative effects on fitness. A single DSB delayed the division to 55 ± 10 min (mean ± SD, n = 765), however, it did not slow down the elongation rate (**Fig. 1f**), and following generations therefore divide faster than average to return to cell size homeostasis. These results show that a single DSB can be repaired remarkably quickly, without fitness cost.

### Pairing between distant homologies

The loss of ParB foci prevented observation of the cleaved *cut-site* locus. However, it has been shown before that markers placed tens of kilobases away from the DSB are protected from RecBCD degradation by correctly oriented *chi* sites^8,9^. Therefore, to visualize the dynamics of the break site, we used a set of *malO*/MalI-Venus (referred to as ‘MalI’) markers integrated at distances *-25kb* or *+170kb* from the *cut-site* (**Fig 2b**). Imaging either of the two MalI markers showed that both sister loci move and pair together in the middle of the cell (**Fig 2a**). The movement of the sister MalI markers was symmetric, unlike previously reported dynamics where the cleaved sister locus moved much further after DSB^10,11^. To test whether the pairing of sister loci is specific, or caused by a global alignment of the chromosomes in response to a DSB, we used a distant MalI marker integrated on the opposite arm of the chromosome (*yahA*, **Fig. 2b**). The distant *yahA* marker maintained its typical positioning during the course of DSB repair (**Fig. 2e, Fig. S5b**), thus excluding a model where HR repair induces pairing of the entire chromosomes. Instead, it appears pairing is specific for the cleaved *cut-site* and its sister locus. Since the sister locus is no different from any other chromosomal locus until it has been located through search, we concluded that the pairing of sister loci implies successful homology search. The *-25kb* markers paired 9.1 ± 3.3 min (mean ± SD, n = 508, **Fig. 2e**) after the DSB, and the homology search is thus faster than this.

**Figure 2.**
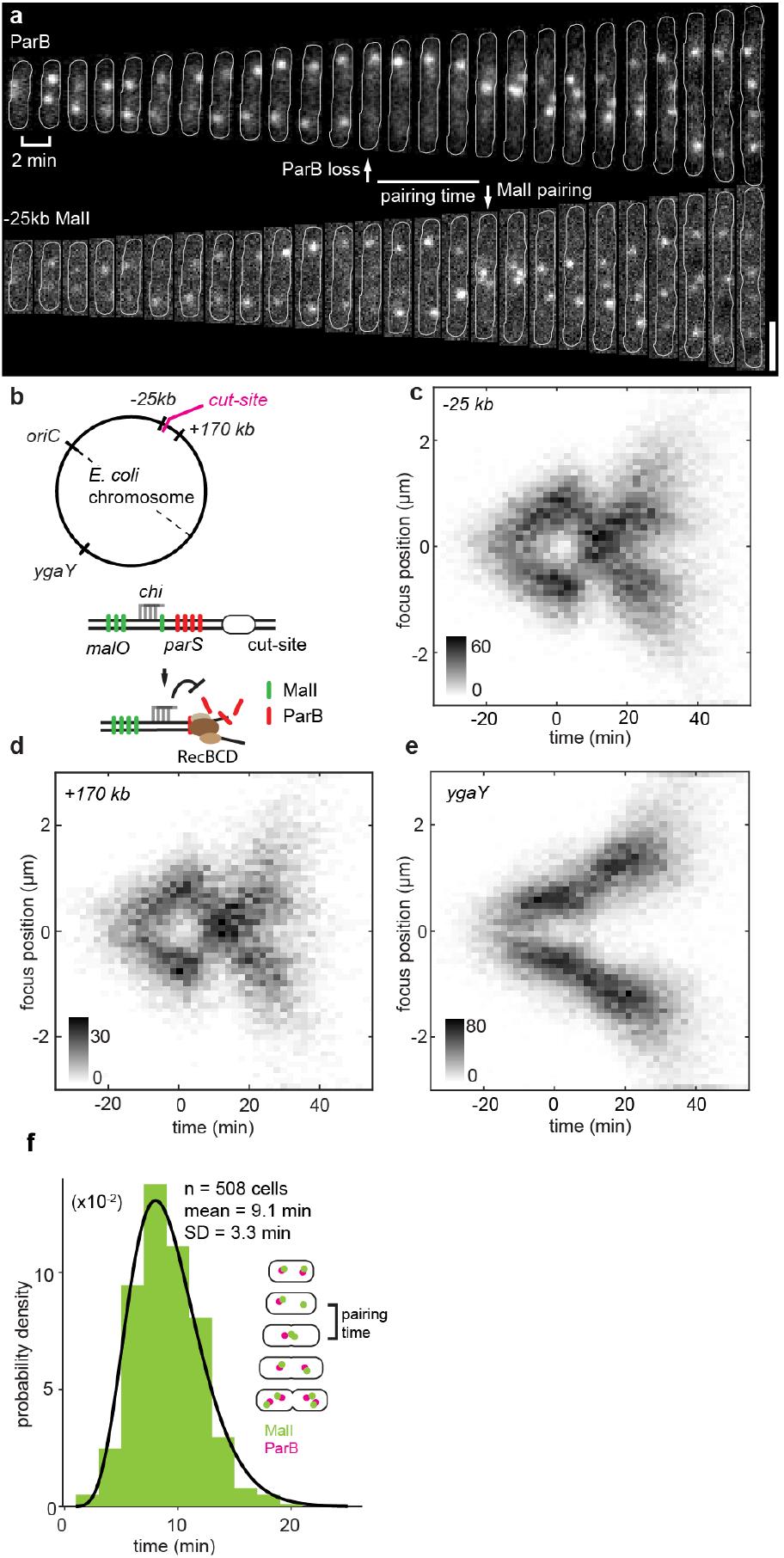
Pairing between segregated sisters. a) A cell with *-25kb* MalI marker undergoing DSB repair. Events of ParB focus loss and pairing of MalI foci are annotated, as well as the pairing time. Cell outlines are displayed with a solid grey line. Scale bar is 2 µm. b) Top: Cartoon showing the circular *E. coli* chromosome with the locations of inserted *cut-site, -25kb, +170kb*, and *ygaY malO*/MalI markers. Bottom: cartoon showing the processing of a DSB in presence of *malO* array flanked by *chi* sites. The *chi* sites prevent RecBCD from degrading the DNA before reaching malO sites, thus *malO* sites are maintained during the repair process. c) Spatial localization of *-25kb* MalI foci in time in cells undergoing DSB. For each cell, the time was aligned based on the time of ParB focus loss. Colorbar displays density of counts. d) Same as in c) but for *+170 kb* MalI foci. e) Same as in c) but for the *ygaY* MalI marker, on the opposite chromosomal arm to the *cut-site*. f) Distribution of MalI foci pairing times for *-25 kb* malI marker during DSB repair. Solid line shows gamma fit to the data (parameters: *k*=8.09, *θ*=1.13).

### RecA filaments are thin and dynamic

The homology search is mediated by a RecA-ssDNA filament^3^. Various structures made by fluorescent RecA fusions have previously been imaged in cells^10–14^. The repair times we measured are, however, much shorter than the lifetimes of the extended RecA structures previously described^10,13^. Therefore, we set out to visualize RecA during DSB repair using a new fluorescent RecA-YFP fusion integrated in tandem with the wild-type *recA* gene in its native operon (**Fig. 3c**). Strains with a similar tandem construct have previously been shown to be fully functional^14^ but have not been used to characterize distant repair events. Our construct likewise retained the wild-type level of DNA-repair activity (**Fig. S3c,g**). We combined the fluorescent RecA construct with the ParB/Cas9 DSB reporter system and studied the repair as described above.

**Figure 3.**
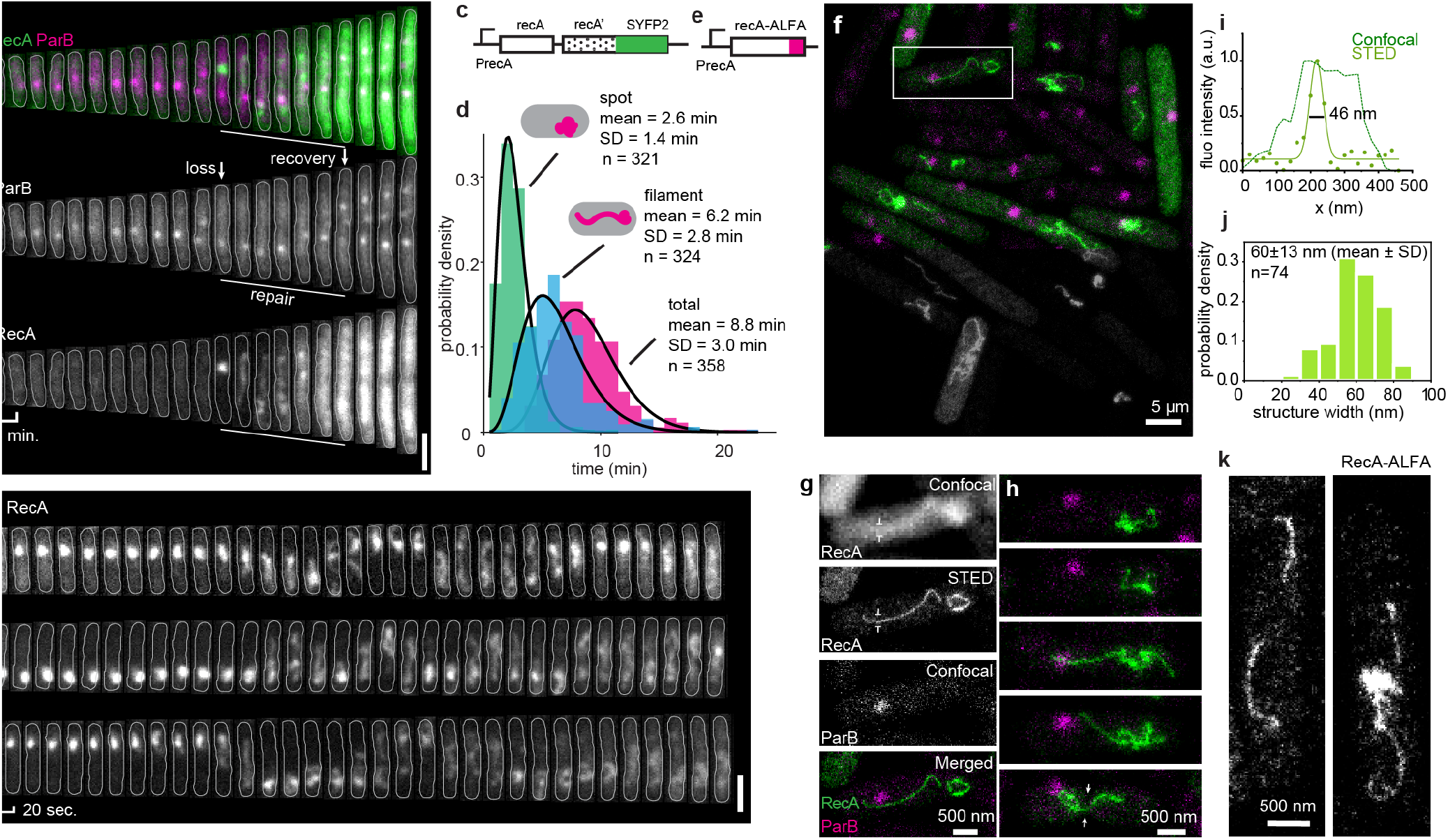
Thin RecA filaments mediate search for homology. a) A cell undergoing DSB repair. Top: False-color composite image of RecA (green) and ParB (magenta) channels. Middle: ParB channel, bottom: RecA channel. Cell outlines are displayed with a solid white line. Scale bar is 2 µm. b) Fast time-lapses of RecA-YFP structures in 3 different cells. Scale bar is 2 µm. c) RecA-RecA-YFP tandem construct inserted into the *recA* locus. d) Lifetimes of RecA structures. Solid line shows gamma fit to the data (parameters: spot: *k*=4.29, *θ*=0.62; filament: *k*=5.28, *θ*=1.18; total: *k*=9.13, *θ*=0.96). e) Cartoon showing the RecA-ALFA construct inserted in the *recA* locus. f) STED image of RecA-ALFA immunolabeled cells. Upper part in false color, superimposed with confocal image of ParB. g) Magnification of the highlighted cell in f), exhibiting an extended RecA structure. Markers indicate where the filament width has been measured. h) Additional examples of cells with either compact or extended RecA-ALFA structures. Note the ongoing septation indicated by arrows. i) Intensity along the filament cross-section labeled in g) (dots) and Gaussian fit to the data (solid line). Filament thickness is measured as full width at half maximum (black solid line). j) Distribution of filament thickness for 74 cells measured as indicated on the left. Mean and standard deviation are indicated. k) STED images of cells harbouring the RecA-RecA-YFP tandem construct immunostained for RecA-YFP.

Induction of DSBs led to the formation of RecA structures at the DSB sites (**Fig. 3a**). Those RecA structures appeared 35 ± 98 sec (mean ± SD, n = 371 **Fig. S3h**) before the loss of ParB focus, and disassembled 6.6 ± 5.2 min (mean ± SD, n = 358 **Fig S3i**) before the repair was finished, as defined by segregation of the ParB foci. We further tested whether those structures were indeed involved in the repair of DSBs. The RecBCD complex is essential for the loading of RecA onto the ssDNA^2^ and consistently, deletion of the *recB* gene abolished the formation of RecA structures after induction of DSBs (**Fig. S4f**). The SOS response is activated by the RecA filament assembled on the ssDNA^3^. We predicted that if the structures we observe are in fact RecA-ssDNA filaments, the life-time of those structures would correlate with the strength of the SOS response. Indeed, the duration of the RecA structure correlated with the signal from the SOS reporter (R=0.36, p=3*10^−12^, **Fig. S3e**). In summary, these results corroborate that the RecA structures we observed after DSBs are in fact RecA-ssDNA.

The RecA structures were wiggly and stretched across the cells at a time scale of tens of seconds. (**Fig 3b**, Supplemental Movies). The persistence length of saturated RecA-ssDNA measured in vitro is ∼900 nm^15^, whereas the in vivo structures appeared to be more flexible. The average lifetime of the RecA structures was 8.8 ± 3.0 min (mean ± SD, n = 358, **Fig. 3d**), and during that time they displayed two forms: (**i**) the initial spot at the DSB location (existed for 2.6 ± 1.4 min (mean ± SD, n = 321)) that increased in intensity (**Fig. S4a**), suggesting RecA loading on ssDNA, and (**ii**) a filament extruding from the initial spot and extending throughout the cell (existed for 6.2 ± 2.8 min (mean ± SD, n = 324)).

It has previously been shown that RecA forms bundle-like structures during DSB repair in *E. coli*^*10*^. To test whether we also could observe such structures in our system, we turned to stimulated emission depletion super-resolution microscopy (STED) to image RecA tagged with the ALFA epitope^16^. The RecA-ALFA alone was fully functional in DSB repair without the need for a wild-type copy (**Fig. S3b,g**). Immunostaining of cells undergoing DSB repair revealed that RecA forms thin, filamentous structures extending throughout the cell (**Fig 3f,g**). Notably, these filaments had uniform thickness, lacking the thick central region characteristic for the previously described bundles^10^. We estimated the thickness of the filaments to be 60 nm (**Fig. 3h**), which is above the 40 nm resolution of the imaging system (**Fig. S4b**). As expected given the high mobility of the RecA-YFP structures in living cells, cells exhibited a large variety of conformations, including complex, tangled up threads, or single winding filaments spanning the length of the cell (**Fig. 3i**). Importantly, we observed the same type of structures when immunostaining RecA-YFP in the tandem construct (**Fig. 3j**), or the RecA-ALFA, RecA-YFP tandem construct (**Fig. S4c-f**). These results show that the RecA structures engaged in fast homology search are thin and remarkably dynamic.

### RecA filament reduces dimensionality of the search

The extended architecture of the RecA-ssDNA filament, and exceptionally fast target search compared to other systems that also rely on homology-directed search^17^, calls for a new quantitative model. Due to slow diffusion rates of both the large RecA-ssDNA complex and chromosomal loci^18^, the search cannot be explained by conventional bi-molecular reaction-diffusion in 3D^19^. To resolve this, we propose that homology search is facilitated by a ‘reduced dimensionality’ mechanism that accelerates the process in two different ways. First, the ATP mediated extension^3^ mechanically stretches the RecA-ssDNA filament across the cell in less than a minute, rapidly covering most of the distance between the broken ends and the search target as shown in **Fig. 3a**. Second, the extended filament interacts with many different sequences in parallel. Such simultaneous probing has previously been suggested based on single-molecule experiments^20^ and cryo-em structures^21^. Our unique addition to the model is the realization that at any z-coordinate (along the cell) there is always at least one segment (the shortest sequence that provides an unique homology) of the stretched RecA-ssDNA filament that is homologous to the dsDNA target sequence (**Fig. 4a**). If we at each point in time consider the search problem from the perspective of the segment of the target sequence that is homologous to the ssDNA-RecA at the current z-coordinate, the complexity is reduced from three to two dimensions. That is, the time of homology pairing is equal to the time it takes for a segment of chromosomal dsDNA to diffuse radially to the RecA-ssDNA filament and not the time it takes for two segments to find each other through 3D diffusion through the whole cell. The difference in time between 3D search and 2D search is in this case approximately 100-fold^22^ (for detailed description see materials and methods).

**Figure 4.**
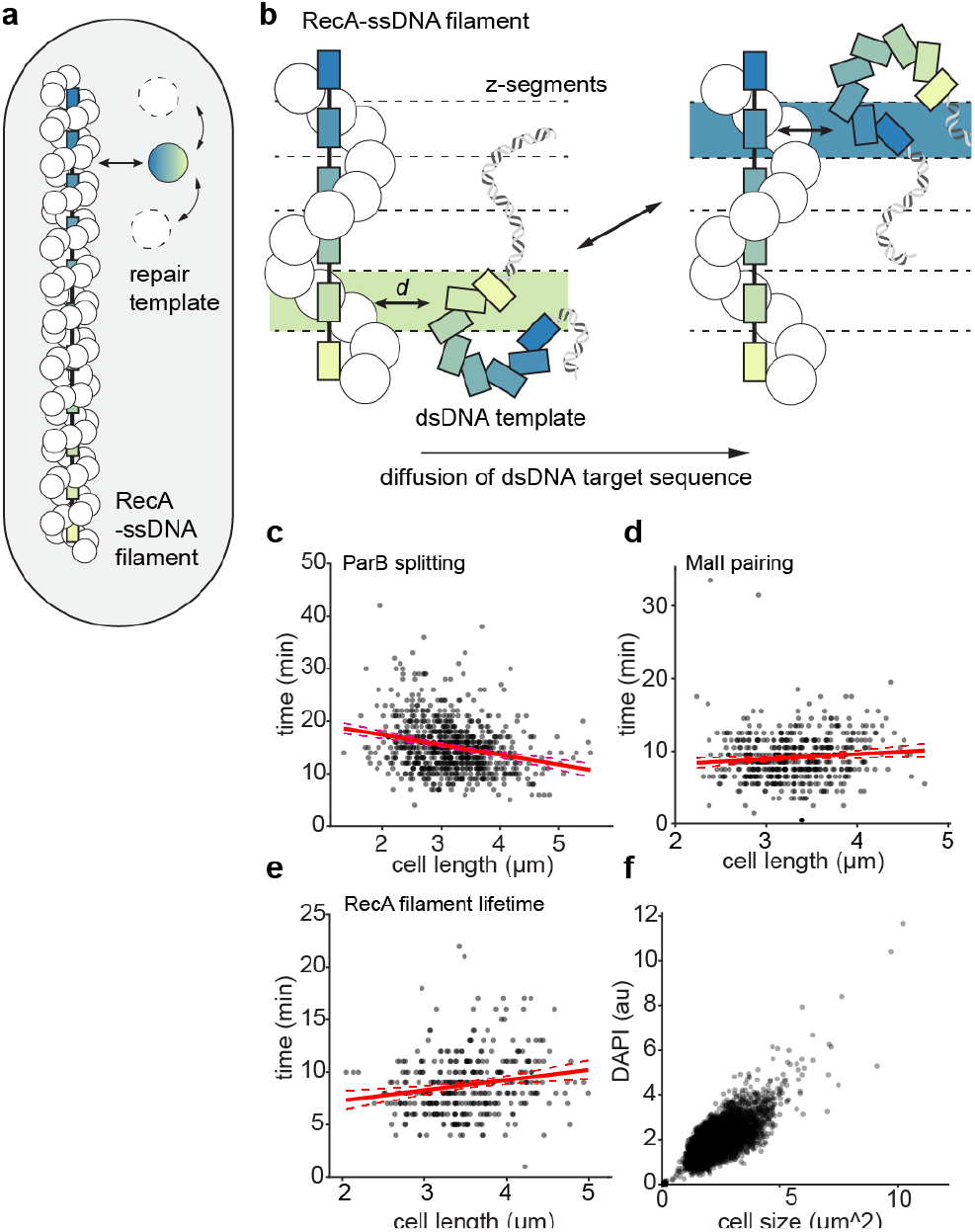
Reduced dimensionality explains fast search times. a) Cartoon showing homology search between extended RecA-ssDNA filament and coiled repair template. The extended filament positions homology to the repair template along the entire cell length b) The RecA-ssDNA filament and the repair template share a homologous segment at each z-coordinate along the length of a cell. Given that during the diffusion of the repair template the homologous segment changes, only the distance to the RecA-ssDNA filament at a given z-coordinate is relevant. The search is now equivalent to the 2D situation, and can be completed within minutes. c) Repair times as a function of cell length at the time of the break. The times were measured based on ParB foci splitting (R = 0.2, p=10^−12^). Red lines display linear fit to the data (solid line), and 95% confidence bounds (dashed lines). d) Search times as a function of cell length at the time of the break. The times were measured based on MalI foci pairing (R = 0.1, p = 0.03). e) Same as in d) but the search times were estimated based on RecA structure life-times (R = 0.2, p=10^−4^). f) DNA amount as a function of cell size as measured by DAPI staining.

In our model, the diffusion of the chromosomal target DNA is as important as that of the RecA filament. The active stretching of the filament is critical to shortening the distance between the homologous sequences to a regime where the diffusive movement of DNA is not prohibitively slow^18^. In the supplementary material, we derive an expression for the expected time for homologous sequences to encounter each other considering the ‘reduced dimensionality’ mechanism. Our model predicts that the search will be completed within 10 min even if each segment of the RecA filament is occupied by probing non-homologous sequences 50% of the time, allowing for an average of 60 ms per probed sequence (see material and methods). Importantly, the model also predicts that the search time should be invariant to the cell length since only the radial distance is relevant. Experimental data confirm that the length of a cell has a minuscule effect on the search time estimated either by ParB foci splitting (**Fig 4c**), RecA structure lifetimes (**Fig 4d**), or MalI foci pairing (**Fig. 4e**), despite that larger cells contain more DNA to explore (**Fig 4f**).

## Discussion

Our findings provide solutions to two key problems in the homology search. First, we show that the stretched RecA-ssDNA filament in a simple and elegant way positions at least one ssDNA segment in the proximity of its homologous sister. Second, since the local search in our model, i.e. after stretching the filament, is driven mainly by the diffusion of the chromosome, it provides a compelling solution to why RecA does not need to hydrolyze ATP in search^23^. ATP-hydrolysis will, however, still be needed for fast dissociation of the RecA-ssDNA filament from the overwhelming amounts of DNA with partial homology^24,25^ that could otherwise entrap the filament. The RecA is a prototypic member of the family of strand exchange proteins, which are found in all forms of life and share a common mechanism^3,5^. We believe that the ‘reduced dimensionality’ mechanism is a conserved property of these proteins. The advantage of the stretched filament is obvious in elongated cells, and long RecA structures were indeed previously observed in *C. crescentus*^*11*^ and *B. subtilis*^*13*^. Interestingly, according to the ‘reduced dimensionality’ model, the search time is not affected by an increase in the amount of DNA as long as the length of the filament scales with the amount of DNA. The mechanism therefore enables search in organisms with genomes larger than bacteria.

Here, we present that search mediated by RecA is remarkably efficient. What was previously considered to be an astonishingly complex step in homologous recombination, is achieved fast and with seemingly no effort. Finally, our robust high-throughput methodology to visualize and dissect DNA repair in real time can be used to address other exciting questions about chromosome dynamics and DNA maintenance.

## Material and Methods

### Strain construction

Strains used in this work are derivatives of E. coli TB28^26^ in which we restored the *rph-1* mutation to wild type and deleted the *malI* gene and maltose operator. Genetic integrations were done using the lambda red integration, resistance cassettes removed using the pCP20 plasmid^27^, and markers were combined using P1 phage transduction.

Labels mCherry-ParBMt1^28^ and MalI-Venus were expressed by constitutive promoters integrated into the *gtrA* locus, respectively. The *malO*/malI marker consists of an array of 12 maltose operators which are the binding site for MalI-Venus.

The DSB cassette consists of the I-SceI recognition site flanked by two lac operators, *parSMt1* site and three chi sites positioned outside the parSMt1 and I-SecI recognition sequence (**Fig. S1**). Cas9 target site was chosen ∼100 bp away from the *parSMt1* site. Notably, the construct is designed in a way that there are no chi sites between the DSB site and *parSMt1* site. The DSB cassette was integrated into *codA* locus.

The RecA-YFP fusion was expressed directly downstream from the endogenous *recA* gene, and made by replacing mCherry with SYFP2 in the construct by Amarh et. al.^14^. The RecA-ALFA fusion was made by introducing ALFA c-terminal to *recA* in either the wild-type *recA* locus or into the tandem construct mentioned above.

List of the strains used in this study can be found in the **Supplementary table S1**.

### Plasmid construction

**p15a-SceIdeg-amp** was cloned using HiFi DNA Assembly (NEB) by fusing two PCR fragments: (1) pSC101SceI_deg_amp^8^ fragment amplified with Jwu035 and Jwu036, and (2) p15aSceI_deg_Kan^8^ fragment amplified with Jwu037 and Jwu038. **p15a-dSceIdeg-amp** was cloned with Gibson assembly from two PCR fragments amplified from p15a-SceIdeg-amp template with two primer pairs: (1) Jwu088 and Jwu090, and (2) Jwu085 and Jwu091. **p15a-Cas9deg-amp** was cloned using Gibson assembly from two fragments: (1) p15a-SceIdeg-amp fragment amplified with Jwu273 and Jwu274, and (2) pCRED (gift from Daniel Camsund) fragment amplified with Jwu263 and Jwu264. Plasmid **p15a-SceIdeg-amp-SOS** was cloned using Gibson assembly protocol with: (1) p15a-SceIdeg-amp fragment amplified with primers Jwu330 and Jwu331, (2) fragment amplified form E. coli genome (from the strain EL1171) with primers Jwu332 and Jwu333. **p15a-Cas9deg-amp-SOS** plasmid was cloned using Gibson assembly with (1) p15a-wtCas9deg-amp fragment amplified with primers Jwu263 and Jwu272 and (2) p15a-SceIdeg-smp-SOS fragment amplified with primers Jwu273 and Jwu274. **p15a-dCas9deg-amp-SOS** was cloned using Gibson assembly protocol using (1) fragment amplified from p15a-SceIdeg-amp-SOS using primers Jwu273 and Jwu274 and (2) fragment amplified from E. coli genome (from the strain EL1605) using primers Jwu264 and Jwu272. **pKD13-recA::ALFA-recAsh-SYFP2-frt-kan-frt** was cloned using golden gate protocol with BbsI restriction enzyme from (1) fragment amplified from *E. coli* genome (from the strain EL2515) with primers Jwu485 and Jwu486, (2) fragment amplified from *E. coli* genome (from the strain EL2699) using primers Jwu487 and Jwu488, and (3) fragment amplified from pKD13-P58-SYFP2-frt-kan-frt with primers Jwu489 and Jwu490.

psgRNA-CS1 was cloned in *E. coli* Top10 by blunt-end ligation of a fragment generated with a PCR with Jwu267 and Jwu184 and psgRNA^29^ plasmid as a template (psgRNA was a gift from David Bikard; Addgene plasmid # 114005), gRNA sequence is ACTGGCTAATGCACCCAGTA.

List of primers used in this study can be found in **Supplementary table S2**.

### High-throughput DSB imaging

#### Growth conditions

For the microfluidic experiments cells were grown at 37°C in M9 media supplemented with 0.4% glucose, 0.08% RPMI 1640 amino acids (Sigma-Aldrich R7131), surfactant Pluronic F-109 (Sigma-Aldrich 542342, 21 µg/ml), and when relevant carbenicillin (40 µg/ml), or kanamycin (20 µg/ml). Cells from -80°C freezer stock were used to inoculate LB media supplemented with adequate antibiotics and grown overnight at 37°C. On the next day the cells were diluted 1/250 in M9 0.4% glucose 0.08% rpmi media and grown for 2 hours, then cells were loaded onto the microfluidic chip. Cells were grown in the microfluidic chip for at least 2 hours before the start of the experiments. Induction of Cas9 and dCas9 was done with a 5 (or 6 minute for spot counting experiments in Fig. S2c) pulse of aTc (20 pg/ml).

#### Microscopy

Microscopy experiments were performed using Ti2-E (Nikon) inverted microscope equipped with CFI Plan Apochromat DM Lambda 100x objective (Nikon), Sona 4.2B-11 sCMOS camera (Andor), and Spectra III (Lumencor) fluorescent light source. The microscope was controlled by Micro-Manager^30^ running in-house build plugins. Fluorescence light source was triggered by the camera with the TTL connection. Fluorescent cubes were used: **CFP** - excitation filter: BrightLine FF02-438/24 (Semrock), emission filter: BrightLine FF01-494/41 (Semrock), dichroic mirror: Di02-R442 (Semrock); **YFP** excitation filter: FF01-514/3-25 (Semrock), emission filter: ET550/50M 200362 (Chroma), dichroic mirror: Di02-R514 (Semrock); **mCherry** excitation filter: FF01-559/34 (Semrock), emission filter: T590LP 262848 (Chroma), dichroic mirror: T585lpxr (Chroma). Imaging was done with 1.5x intermediate magnification lens. Phase contrast images were taken with the CFP cube inserted. Typically, a phase contrast image was acquired every minute with 80 ms exposure, CFP channel every third minute with fluorescence light intensity set at 5% and exposure of 40 ms, mCherry channel was acquired every minute with fluorescent light intensity set at 20% and exposure of 80 ms, YFP channel was acquired every minute (for imaging RecA), or every second minute (for MalI experiments) with fluorescence light intensity set at 40% and exposure of 100 ms. For spot counting experiments in **Fig. S2c** the mCherry channel was imaged every 3rd minute, and 3 z-slices separated by 300 nm were taken at every time point and fluorescent images were reconstructed using maximum intensity projection.

#### Microfluidic experiments

The microfluidics experiments were performed using PDMS mother machine microfluidic chip developed previously^7^. This chip design allows for loading of two different strains and for automated switching of the media. Media pressure was controlled using OB1 MK3+ microfluidic flow controller (Elveflow). The aTc was pulsed at the beginning of the for 3h experiments, or 20 minutes after the start of image acquisition for 4h experiments.

#### Chromosome staining with DAPI

Strain EL2504 and EL1743 (with *dnaC2*^*31*^) were grown overnight in LB media. Next day, cell cultures were diluted in 5 ml of M9 with 0.4% glucose and 0.08% RPMI 1640 amino acids (Sigma-Aldrich R7131), and incubated in 37°C (strain EL2504). EL1734 strain was grown at 30°C and 2 h before imaging one aliquot of the culture was grown at 42°C to induce the replication arrest. Then DAPI was added to the final concentration of 1 µg/ml and cells with DAPI were incubated for 30 minutes at the growth temperatures. Then 1 ml of cells was centrifuged at 4°C, 7000 RPM (5424 R, Eppendorf) for 3 minutes, resuspended in 50 ul of cold ITDE (IDT) buffer supplemented with 10 mM MgCl2. 2 ul of concentrated cells was mounted on an agarose pad for microscopy. Imaging was done with the 445 laser with the power 12 mW/cm^2^, and exposure time of 220 ms. Phase contrast images were segmented using Nested-Unet neural network^32^, trained in-house. DAPI images were corrected for background by subtracting the mean pixel intensity of the area outside of the cell.

#### Image analysis

Data analysis was done in MATLAB (Mathworks), with the exception of the segmentation that was done in Python. Microscopy data was processed using an automated analysis pipeline developed previously^33^, however with several modifications. First, segmentation of phase contrast images was done using Nested-Unet neural network^32^ trained in-house, specifically for our microscopy setup. We trained two networks, one to segment cells, and another to detect growth channels on the phase contrast images. Transformation matrices between images acquired with different filter cubes were measured and fluorescent images were transformed to correct for the pixel shifts between fluorescence and phase contrast images. Gramm toolbox was used to generate some of the plots in Matlab (https://github.com/piermorel/gramm)

#### Selection and analysis of cells undergoing DSB

Cells undergoing a DSB repair were selected based on at least 4-fold increase of CFP signal from plasmid borne SOS-reporter. First, the CFP image was background corrected by subtracting an image filtered with a Gaussian filter with the kernel size of 20 pixels (using matlab function ‘imgaussfilt’) from the original CFP image. We limited the analysis to the cells that had no major errors in segmentation, lived for at least 9 minutes, and divided during the experiment. Manual repair dynamics annotation was done only on cells that contained two ParB foci prior to the DSB, and divided after the repair. Cells that had more than two ParB foci, or that induced a DSB more than once were excluded from the analysis. In the experiment in Fig. S2c ParB foci were detected automatically using radial symmetry-based method. Spot positions were mapped on the cell length using a custom written Matlab code.

### Superresolution STED microscopy

#### Sample preparation

Cells expressing RecA-ALFA, RecA-YFP or both, were grown for 3 hours at 37°C in M9 media with 0.4% glucose, RPMI 1640 amino acids (0.08%), carbenicillin (20 µg/ml) and kanamycin (10 µg/ml). Then, Cas9 was induced for 40 min with aTc (0.8 pg/ml), after which the cells were fixed with formaldehyde (3.5%) for 10 min. Fixing was quenched with 100 mM glycine and the cells washed in PBS before permeabilization in 70% ethanol for one hour. For staining, the cells were blocked in PBS with BSA (1%) for 30 minutes and then incubated with antibodies at 1:200 dilution for at least one hour. We used camelid single domain antibodies conjugated to either Star635P or Star580, for RecA-ALFA FluoTag-X2 anti-ALFA (N1502-Ab635P) and for RecA-YFP FluoTag-X4 anti-GFP (N0304-Ab635P and N0304-Ab580, NanoTag Biotechnologies). After three washes in PBS, the cells were mounted on an agarose pad for microscopy.

#### STED microscopy

The super-resolution imaging was performed with a custom-built STED setup^34^. Excitation of the dyes was done with two pulsed diode lasers, one at 561 nm (PDL561, Abberior Instruments) and one at 640 nm (LDH-D-C-640, PicoQuant), which were coupled into a fibre in order to co-align the beams and shape the wave fronts. A beam at 775 nm was used as a depletion laser (KATANA 08 HP, OneFive). The depletion beam was shaped to a donut in the focal plane using a spatial light modulator (LCOS-SLM X10468-02, Hamamatsu Photonics). The laser beams were focused onto the sample using a HC PL APO 100×/1.40 Oil STED White objective (15506378, Leica Microsystems), through which also the fluorescence signal was collected. The imaging was done with a 561 nm excitation laser power of 8-20 µW, a 640 nm excitation laser power of 4-10 µW and a 775 nm depletion laser power of 128 mW, measured at the first conjugate back focal plane of the objective. 2-color STED imaging of RecA-YFP together with RecA-ALFA was done in a line by line scanning modality, averaging over 4 or 8 lines; while ParB and RecA-ALFA was recorded frame by frame, with the first channel in confocal and second channel in STED. The pixel size for all images was set to 20 nm with a pixel dwell time of 50 µs. Raw images were processed and visualized using the ImSpector software (Max-Planck Innovation, Goettingen, Germany) or ImageJ (https://fiji.sc/). Brightness and contrast were linearly adjusted for the entire images. The size of the RecA filaments was calculated tracing line profiles perpendicular to the filament orientation and averaged on 2 pixels on the raw images. The data were then fitted with a Gaussian function with the software OriginPro2020, from which the full width half maximum was extracted. The resolution of the microscope was measured on a calibration sample, made of sparse antibodies attached to the glass, coupled with the STAR635P dye. The line profiles were extracted and fitted with a Gaussian function, from which the full width at half maximum was extracted as the dot size.

### I-SceI enzyme experiments

For experiments with the I-SceI meganuclease, cells with the p15a-ISceI plasmid were cultured in M9 minimal medium, loaded in a microfluidic chip and then incubated at 37°C using M9 medium with glucose (0.4 %), RPMI 1640 amino acid supplement (Sigma-Aldrich R7131, 0.05 %), carbenicillin (20 µg/ml) and Pluronic F-108 (21 µg/ml). DSBs were induced by switching for three minutes to media also containing aTc (20 ng/ml) and IPTG (1 mM), and then for three minutes to media with only IPTG. The cells were then imaged while repairing and recovering in the initial medium.

### Serial-dilution plating assay

Bacteria cultures were streaked on LA plates from freezer stock and grown overnight at 37°C. Following day, 5 ml of LB media was inoculated with a single colony from the overnight plate and cultured for 6 h at 37 degrees. Next, 10-fold serial dilutions were made in LB and 4 μl of each dilution was plated on a LA plate, or LA plate containing 1 μg/ml of Nalidixic acid. Plates were incubated overnight at 37°C.

## Search model by extended RecA filament

Homology search will be treated as a diffusion limited reaction with a transport time for the homologous sequences to reach a reaction radius and a probing time once the sequences have met. The probing time for the correct sequence will be negligible compared to the time for getting the homologous sequences in contact, but the overall reaction will be slowed down by the overwhelming number of incorrect sentences that will need to be probed.

To quantify the situation, we will start with few approximations. Assume that the RecA-ssDNA filament is a thin rod in the center of the cylindrical nucleoid reaching from pole to pole in the z-direction, whereas the homologous dsDNA sequence is coiled up at a random position in the nucleoid. The relevant time for the homologous sequences to find each other is the time for a segment in the coiled up dsDNA to diffuse radially into the rod in the center of the cell. The central realization in our model is that it does not matter at which z-coordinate it reaches the rod. This transforms the search problem from 3D to 2D, since we can describe the search process from the perspective of the dsDNA fragment that is homologous to the ssDNA sequence that happens to be at the z-position where the rod is reached first.

An equivalent, but slightly more stringent, way to think about the situation, is to consider that the first binding event for many independent searchers (i.e. the chromosomal dsDNA segments), that each can bind to one out of many targets (i.e. the RecA-bound ssDNA segments), has the same rate as one searcher that can bind all targets. The total rate of binding is *r* = ∑_*i*_ *r*_*i*_, where *r*_*i*_ is the rate for template segment to *i* find its homologous ssDNA segment. If we write out the dependence of at which z-coordinate, *z*_*j*_, the filament is reached, the total rate can be expressed as *r* = ∑_*j*_ ∑_*i*_ *r*_*i*_(*z*_*j*_)*p*(*z*_*j*_), where *p*(*z*_*j*_) is the probability to reach the filament at position *z*_*j*_ and *r*_*i*_(*z*_*j*_) is the conditional rate for segment *i* of binding given that the filament is reached at *z*_*j*_. Here, the rate of binding is zero unless the template segment matches the ssDNA that is at the specific z-position, i.e. *r*_*i*_(*z*_*j*≢*i*_) = 0 which means that *r* = ∑_*i*_ *r*_*i*_(*z*_*i*_)*p*(*z*_*i*_), i.e. the total binding rate is the same as the binding rate for a single dsDNA segment that can bind at any position at the filament, irrespective of z position, and for which each position is homologous.

### Search time prediction for *E. col*i

In a first order approximation of how long it takes for a chromosomal dsDNA segment to diffuse from a random radial position in the nucleoid to the filament in its centre, we can use the diffusion limited rate for reaching a rod in the centre of a cylinder of length 2L, i.e. k = 2π (2L) D / ln(R/r)^22^, where r is the reaction radius of the rod, which is assumed to be in the order of the size one nucleotide (1nm), and R is the nucleoid radius. The concentration of the searching dsDNA fragment is c = 1/V = 1/(2LπR^2^). The average time to reach the rod is therefore T = V/k = R^2^ln(R/r)/2D. The nucleoid radius R ≈ 200 nm and the reaction radius is in the order of r ≈ 1 nm, although the exact value is inconsequential since only its logarithm enters into the time. The complicated parameter is the diffusion rate constant D, since DNA loci movement is sub-diffusive and D is therefore lower at a long length scale than a short. The process will however be limited by the long distance movement corresponding to R. At the length scale of R = 200 nm, D_R_ ≈ 0.0007 μm^2^s^-1 18^. The association step of the search process is thus T ≈ (0.2 μm)^2^ln(200 nm/1 nm)/0.0007 μm^2^s^-1^ = 300 s ≈ 5 min. If we consider that also the RecA filament is moving on the minute timescale this will only speed up search further.

### Time for probing

The RecA filament will however not be accessible for binding all the time since it also needs to probe all other dsDNA segments. If only half is needed for homologous sequences to meet, 5 min is still available for probing other sequences. Is this sufficient to probe all sequences? If the dsDNA is probed in n bp long segments, each of the equally long segments of the 2 kb long ssDNA in the RecA filament will have to interrogate, on average, every 2000/n segment of the chromosome. There are 9.6 Mbps (4.8 Mbps / genome x ∼2 genomes / cell) of dsDNA in the cell, which corresponds to 9.6×10^6 / n probing segments. This, in turn, means that each ssDNA segment in the RecA filament needs to test (9.6×10^6 / n) / (2000 / n) = 4800 dsDNA segments. The average time for each test cannot be longer than 300 s / 4800 ≈ 63 ms which should be plenty of time considering that Cas9-sgRNA takes at average 30ms to make a similar task^17^.

### Alternative models

Alternative models for how to get sequences sufficiently close to probe for homology can come in many other flavours.

The most naive comparison is considering the bi-molecular reaction of a “particle” with diffusion rate corresponding to the dsDNA segment and a diffusion limited binding to a non-diffusive segment of the filament with a reaction radius corresponding to r above (1 nm). Here, the rate of the diffusion limited reaction is k=4πrD^35^, whereas the concentration of segment is the same as above (c = 1/V = 1/(2LπR^2^)). This results in T = V/k = LR^2^/2rD_L_, which should be compared to R^2^ln(R/r)/2D_R_ in the 2D situation. Importantly D_L_, the diffusion rate at the length scale of the cell is one order of magnitude lower than D_R_^18^. The ratio is (L/r) / ln(R/r)≈175, (D_R_/D_L_)≈1750 considering L=1um, R=300nm and r=1nm. D_R_/D_L_≈10. The actual value for r is more important in this case, because the number of rebinding events is more important than in the 2D case.

An intermediate step towards the 2D model is to parallelize the naive model and think of the homologous dsDNA as divided in segments that can search in parallel and independently for their respective homology in the RecA filament. For example, if we divide a 1750 bases long filament into segments of 10 there we would get a speed up of 175 times and the remaining difference compared to the 2D model would be (D_R_/D_L_)-fold difference of short and long range diffusion.

A final model is to consider the homologous DNA as static and that only the RecA filament moves like a knitting needle in a ball of yarn. This situation is not as straightforward to quantify, since we do not know how rigid the RecA filament is on different length scales in the cell. It appears flexilel on the 100nm length scale, but the probing interactions will have to occur on the 1 nm scale and we therefore have no knowledge of how fast the filament would explore the genome. It can clearly probe many DNA segments simultaneously^20^ but it causes complex constraints to probe sequences with one part of the filament at the same time as the filament should move to explore the rest of the genome. Detailed simulations may be needed to predict the expected search times for this type of model.

## Acknowledgements

We wish to thank David Fange, Iremli Barkefors and Daniel Jones for helpful input on the manuscript, Daniel Camsund for the pCRED plasmid, Spartak Zirkin and Praneeth Karempudi for help with the image analysis. J.W is an EMBO non-stipendiary fellow (EMBO ALTF 1029-2018). This work was funded by the H2020 Marie Curie Individual Fellowship grant, Agreement Number 842047 for J.W., European Research Council (ERC), the Swedish Research Council (VR), and the Knut and Alice Wallenberg Foundation (KAW).

## Author contributions

J.E., J.W. and A.H.G. conceived the study and interpreted results. A.H.G., J.W., J.L. and P.L. conducted the experiments. J.W. & A.H.G. wrote analysis code and analysed data. A.H.G., J.W. and P.L. constructed the strains, J.E. derived equations. G.C. and I.T. contributed the STED component. J.E., A.H.G. and J.W. wrote the manuscript with the assistance of all authors.

## Code availability

The computational code that supports the findings of this study are available from the corresponding author upon reasonable request.

## Data availability

The data that support the findings of this study are available from the corresponding author upon reasonable request.

---

**Figure S1.**
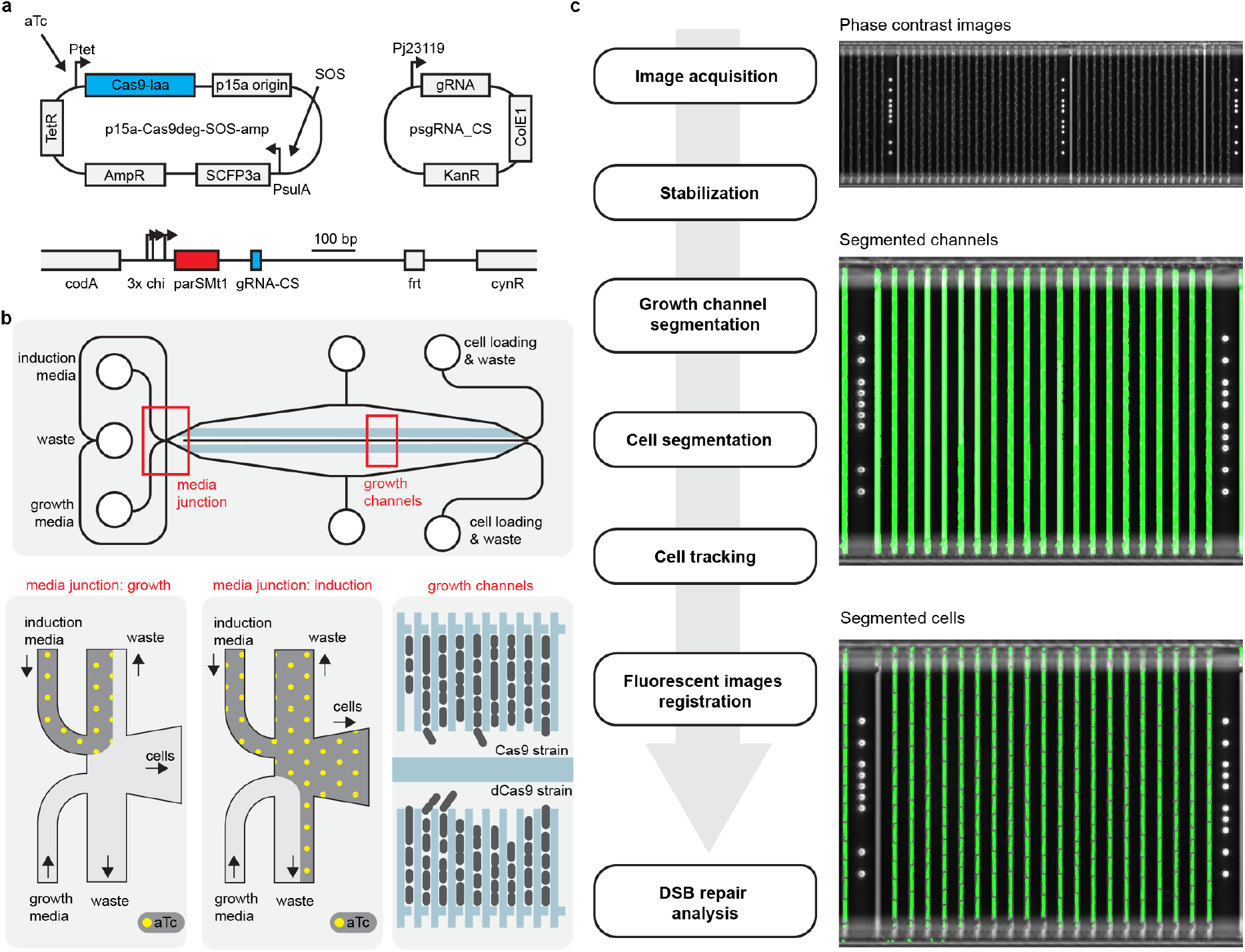
DSB induction system and image processing. a) Top: Plasmids used in the DSB reporter system. Cas9 and SOS-reporter are encoded on a single plasmid, sgRNA on a separate plasmid. Bottom: *cut-site* cassette integrated into *codA* locus on the chromosome. b) Top: Cartoon showing a simplified schematic of microfluidic chip. Bottom: cartoons showing a media input junction and flow of the media during growth (left) and induction (right) phases of the experiment. The rightmost cartoon shows an arrangement of the growth channels on the chip. Typically, strains transformed with an active Cas9 and dCas9 were loaded on two sides of the chip, to serve as a control for induction strength. c) Left: Steps in the image analysis pipeline. Right: *Top* image shows a full field of view (FOV) of a single position of the microfluidic chip. Typically, 16 positions were imaged during an experiment. *Middle* image shows an enlarged section of the FOV with an overlaid mask of segmented channels. *Bottom* image shows the same section of FOV as in the middle, but overlaid are segmented cells.

**Figure S2.**
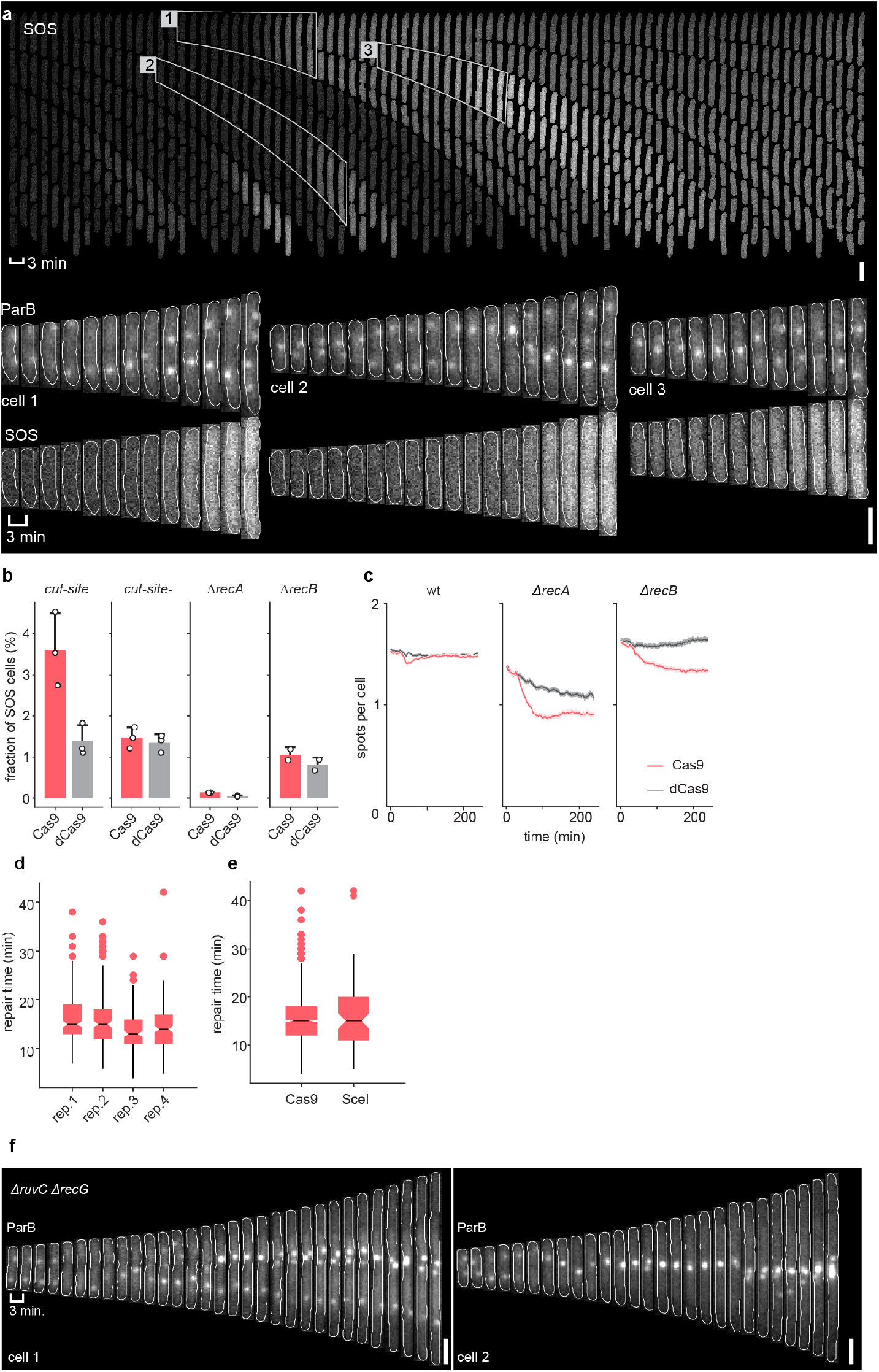
Specificity of the DSB induction. a) Top: Cells during a DSB experiment showing SOS reporter channel, cells that activated SOS response are outlined and displayed on the bottom. Scale bar is 2 µm. b) Fractions of cells that activated SOS response in different strains and for active, or dead Cas9 variants. Cut-site-strain lacks chromosomal *cut-site*. Bars represent the mean of at least 2 experiments, error-bars show standard deviation. Dots represent a mean for each of the independent experiments. c) Mean ParB foci number per cell after induction of Cas9 or dCas9 (induction was at t = 20 min). Solid line shows mean, light coloured area shows 95% confidence interval measured by bootstrap. d) Repair times measured by ParB focus loss to recovery in each of 4 experimental replicates. Centerline displays the median, bottom and top edges show 25th and 75th percentiles, whiskers show most extreme data points not considering outliers, and the solid dots show outliers. e) Repair times measured by ParB focus loss to recovery of DSBs caused by either Cas9 or I-SceI enzyme. Centerline displays the median, bottom and top edges show 25th and 75th percentiles, whiskers show most extreme data points not considering outliers, and the solid dots show outliers. f) *∆ruvC ∆recG* cells after DSB. Scale bar is 2 µm.

**Figure S3.**
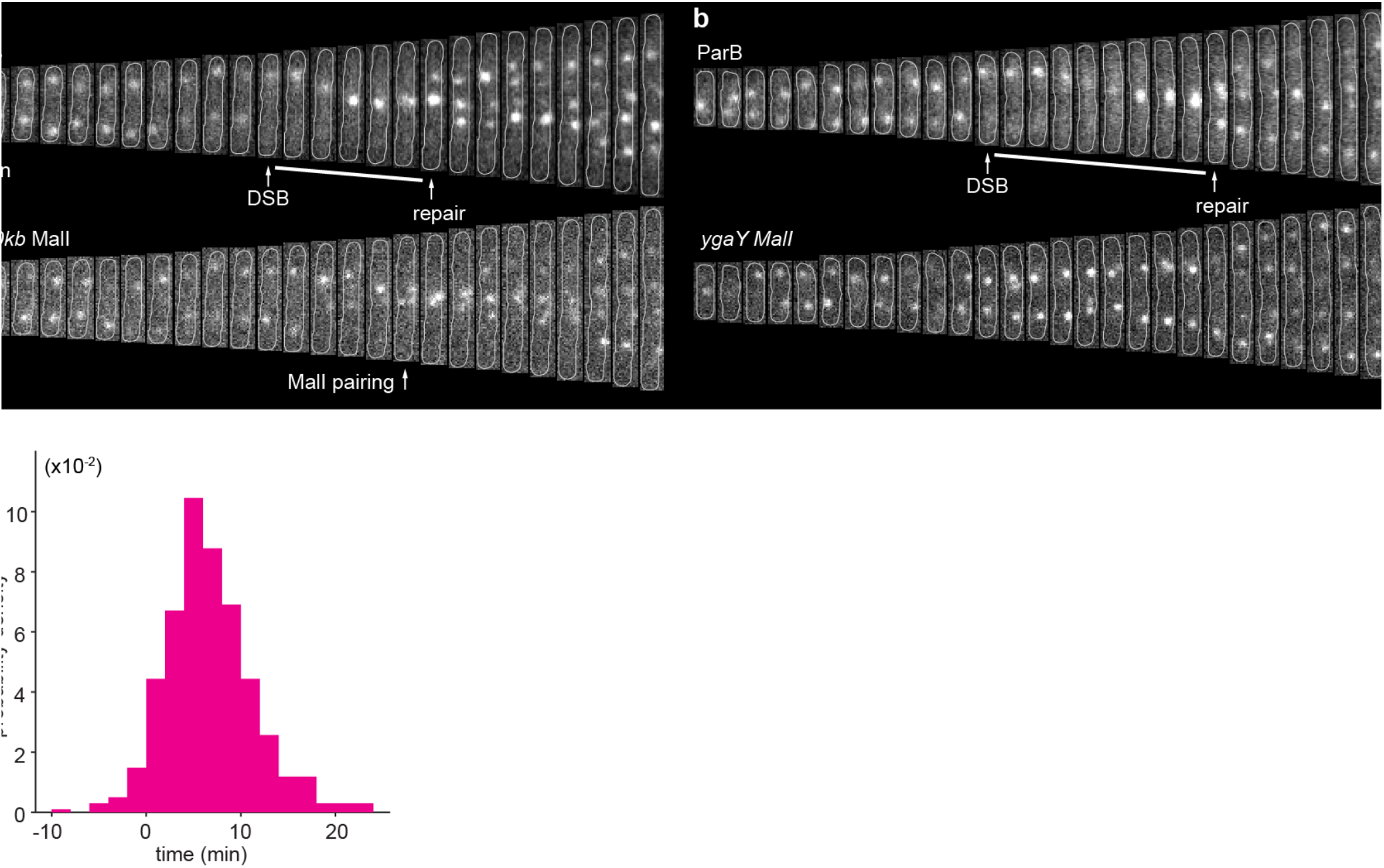
Paring of the distant homologies. a) Cell with *+170kb* MalI marker undergoing DSB repair. ParB focus loss and recovery events are shown. Scale bar is 2 µm. b) As in a), but for a acell with *ygaY* MalI marker. Event of MalI markers pairing is shown Scale bar is 2 µm. c) Time of ParB focus splitting after pairing of the *-25 kb* MalI foci.

**Figure S4.**
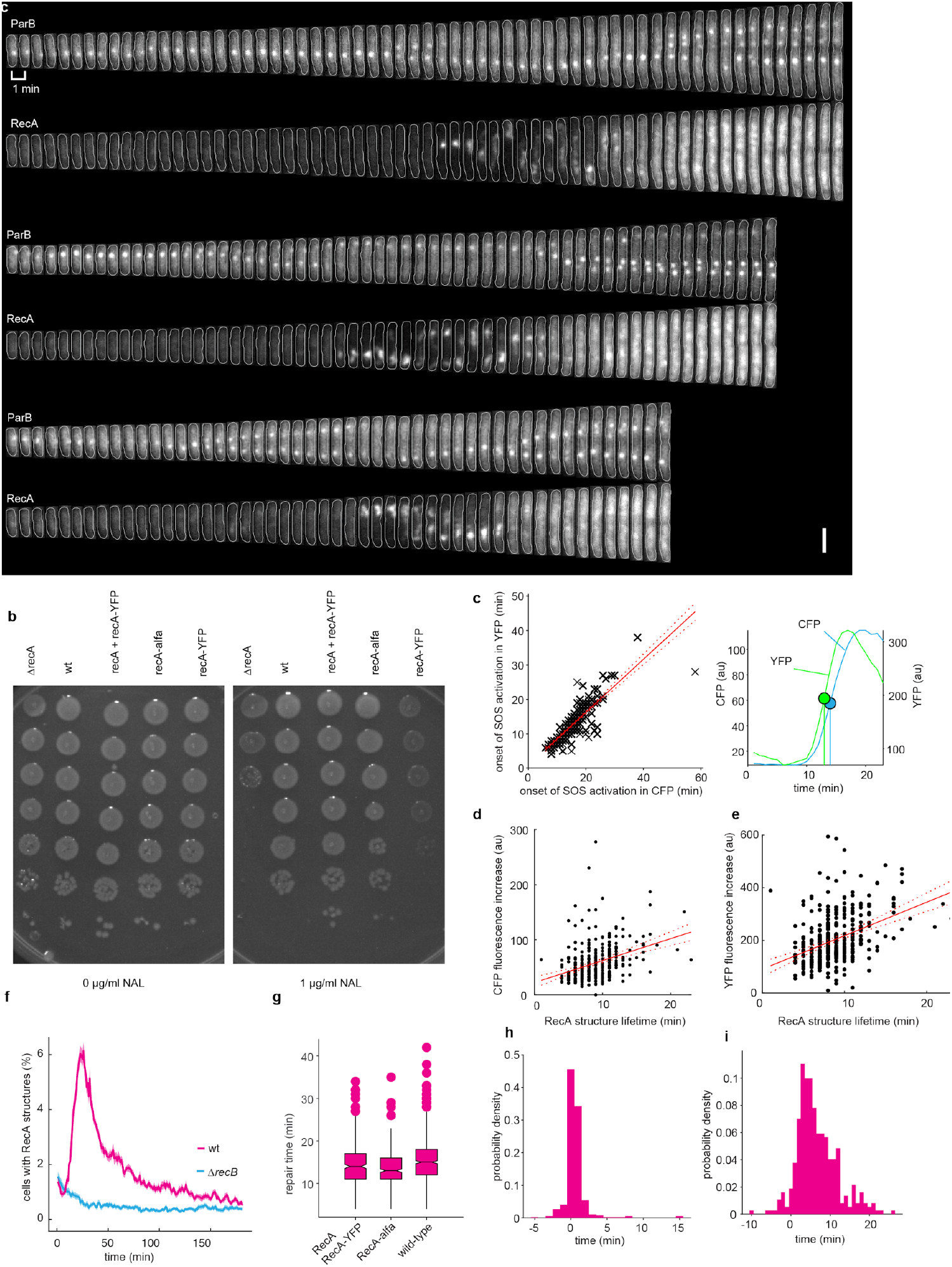
RecA structures are specific. a) Cells with ParB and RecA labels undergoing DSB repair. Scale bar is 2 µm. b) Growth of strains with different RecA variants without, or in presence of DNA damage induced by Nalidixic acid. c) Correlation between the time of activation of SOS response measured by fluorescence from plasmid SOS reporter and from RecA expression. SOS activation was measured by the time at which fluorescent signals reached half maximum value. Analysis was restricted only to cells that activated SOS response. Red line shows linear fit to the data (black), dashed lines show confidence bounds. Right-hand plot shows SOS (blue) and RecA (green) intensity traces for a single cell. Time of half-maximum signal for each channel is shown. d) Correlation between the lifetime of RecA structure and increase of SOS signal intensity for cells undergoing DSB repair. Solid red line shows linear fit, dashed red lines show confidence bounds. e) Same as in d) but for a correlation between RecA structure lifetime and increase in expression of the RecA-YFP. f) Fraction of cells with RecA structure after the induction of DSB (at time=0). g) Repair times measured by ParB focus loss to recovery of DSBs for different RecA variants. Centerline displays the median, bottom and top edges show 25th and 75th percentiles, whiskers show most extreme data points not considering outliers, and the solid dots show outliers. h) Times between appearance of RecA structure to loss of ParB focus. i) Times between the disassembly of RecA structure to splitting of ParB focus.

**Figure S5.**
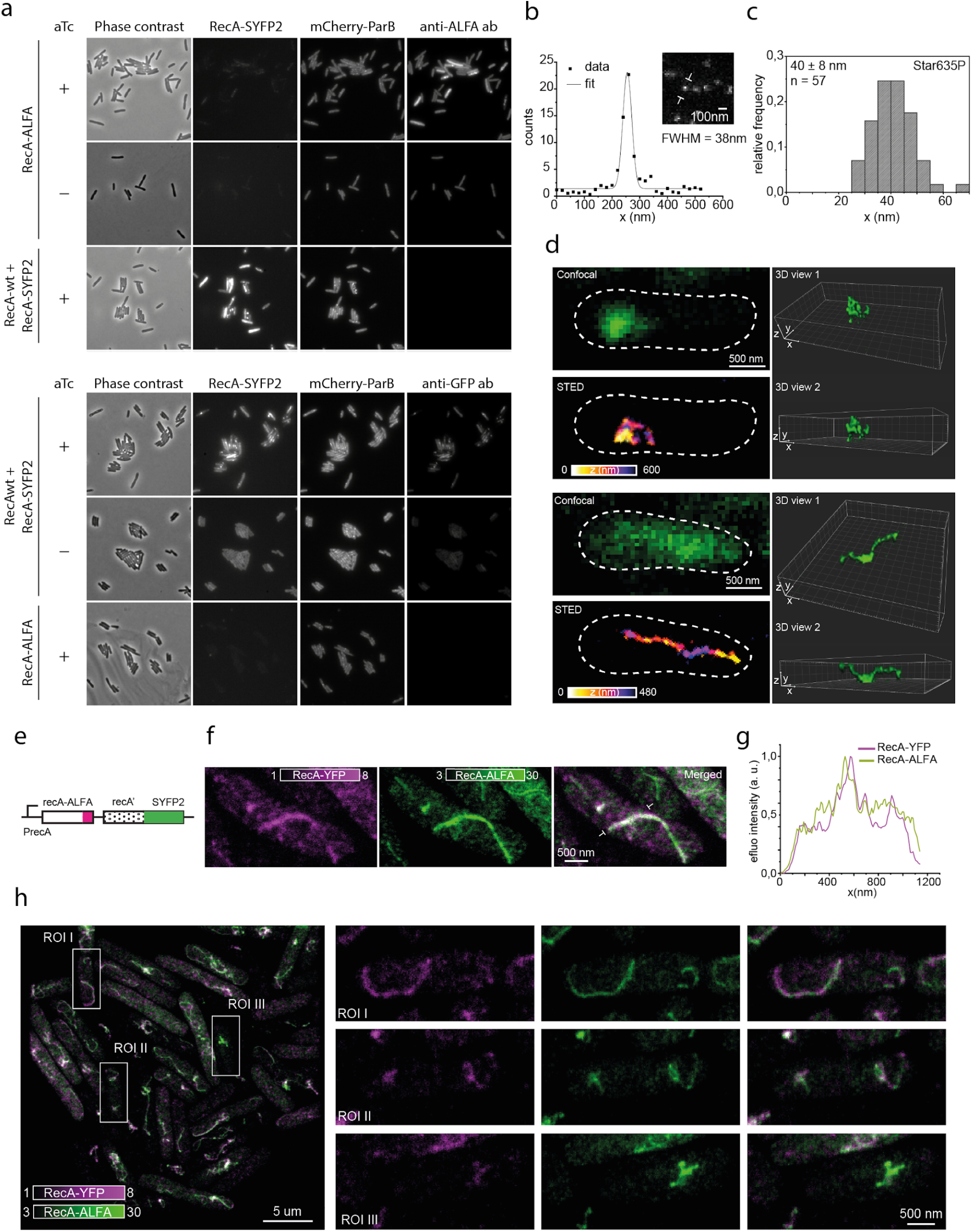
Validation of RecA immunostaining. a) Specificity of intracellular labelling of RecA-ALFA and RecA-YFP using camelid single domain antibodies. b) Measurement of STED resolution using Star635P-conjugated antibodies: Profile over single antibody as indicated in the insert and Gaussian fit. c) Distribution of full width at half maximum of Gaussian fitted line profiles of antibodies, representing the resolution of the STED microscope. d) Examples of three-dimensional STED images of immunostained RecA-ALFA structures *recA-alfa-recA-syfp2* construct inserted into the *recA* locus e) Genetic construct expressing both RecA-ALFA and RecA-YFP, inserted in the native *recA* locus. f) STED images of with extended RecA filament, stained with anti-GFP-Star580 and anti-ALFA-Star635P antibodies, imaged in their respective channels. h) Fluorescence intensity profile as indicated in e. i) Further STED image with immunolabeled cells as in f. Magnifications of the labelled regions. (right)

**Table S1.** Strains used in this study

| strain name | genotype |
| --- | --- |
| EL666 | ΔrecA::frt-kan-frt |
| EL1171 | Eco MG1655 rph+ ΔmalI::frt intC::P59-malIvenus-frt gtrA::P58-mCherry:parB-SpR codA::lacOsym-lacOCS-parBMtl-frt yahA::mal012x-frt yibU::Psula'-scfp3a-KanR |
| EL2504 | Eco MG1655 rph+ lac+ ΔmalI::frt gtrA::P58-mCherry:parB-SpR codA::parBMtl-lacOsym-lacOCS-frt |
| EL2515 | Eco MG1655 rph+ lac+ ΔmalI::frt gtrA::P58-mCherry:parB-SpR codA::parBMtl-lacOsym-lacOCS-frt recA-recX::recAmwg-SYFP2-frt |
| EL2561 | Eco MG1655 rph+ lac+ ΔmalI::frt gtrA::P58-mCherry:parB-SpR codA::parBMtl-lacOsym-lacOCS-frt / p15a-Cas9deg-SOS-amp + psgRNA-CS1 |
| EL2562 | Eco MG1655 rph+ lac+ ΔmalI::frt gtrA::P58-mCherry:parB-SpR codA::parBMtl-lacOsym-lacOCS-frt / p15a-dCas9deg-SOS-amp + psgRNA-CS1 |
| EL2618 | Eco MG1655 rph+ lac+ ΔmalI::frt codA::parBMtl-CS1null-lacOsym-lacOCS-frt / p15a-wtCas9-SOS-amp psgRNA_CS1 |
| EL2619 | Eco MG1655 rph+ lac+ ΔmalI::frt codA::parBMtl-CS1null-lacOsym-lacOCS-frt / p15a-dCas9-SOS-amp psgRNA_CS1 |
| EL2621 | Eco MG1655 rph+ lac+ ΔmalI::frt gtrA::P58-mCherry:parB-SpR codA::parBMtl-lacOsym-lacOCS-frt recA-recX::recAmwg-SYFP2-frt / p15a-wtCas9deg-SOS-amp + psgRNA_CS1 |
| EL2622 | Eco MG1655 rph+ lac+ ΔmalI::frt gtrA::P58-mCherry:parB-SpR codA::parBMtl-lacOsym-lacOCS-frt recA-recX::recAmwg-SYFP2-frt / p15a-dCas9deg-SOS-amp + psgRNA_CS1 |
| EL2653 | Eco MG1655 rph+ ΔmalI::frt gtrA::P58-mCherry:parB-SpR codA::parBMtl-lacOsym-lacOCS-frt ΔrecG::frt ΔruvC::frt/ p15a-wtCas9deg-SOS-amp + psgRNA_CS1 |
| EL2672 | Eco MG1655 rph+ lac+ ΔmalI::frt gtrA::P58-mCherry:parB-SpR codA::parBMtl-lacOsym-lacOCS-frt ΔrecA::frt / p15a-wtCas9-deg-amp + psgRNA_CS1 |
| EL2673 | Eco MG1655 rph+ lac+ ΔmalI::frt gtrA::P58-mCherry:parB-SpR codA::parBMtl-lacOsym-lacOCS-frt ΔrecA::frt / p15a-dCas9-deg-amp + psgRNA_CS1 |
| EL2674 | Eco MG1655 rph+ lac+ ΔmalI::frt gtrA::P58-mCherry:parB-SpR codA::parBMtl-lacOsym-lacOCS-frt ΔrecB::frt / p15a-wtCas9-deg-amp |
|  | + psgRNA_CS1 |
| EL2675 | Eco MG1655 rph+ lac+ ΔmallI::frt gtrA::P58-mCherry:parB-SpR<br>codA::parBMt1-lacOsym-lacOCS-frt ΔrecB::frt / p15a-dCas9-deg-amp<br>+ psgRNA_CS1 |
| EL2699 | Eco MG1655 rph+ lac+ ΔmallI::frt gtrA::P58-mCherry:parB-SpR<br>codA::parBMt1-lacOsym-lacOCS-frt recA:ALFA-frt-KmR-frt |
| EL2801 | Eco MG1655 rph+ lac+ ΔmallI::frt codA::parBMt1-lacOsym-lacOCS-frt<br>intC::P59-mallIvenus-frt gtrA::P58-mCherry:parB-SpR yahA::mal012-<br>frt-cmr-frt / p15a-wtCas9-SOS psgRNA_CS1 |
| EL2802 | Eco MG1655 rph+ lac+ ΔmallI::frt codA::parBMt1-lacOsym-lacOCS-frt<br>intC::P59-mallIvenus-frt gtrA::P58-mCherry:parB-SpR ygaY::mal012-<br>frt-cmr-frt / p15a-wtCas9-SOS psgRNA_CS1 |
| EL2865 | Eco MG1655 rph+ lac+ ΔmallI::frt gtrA::P58-mCherry:parB-SpR<br>codA::parBMt1-lacOsym-lacOCS-frt ybbD::mal012x-frt-Cmr-frt /<br>p15a-wtCas9deg-sos-amp / psgRNA_cs1 |
| EL2834 | Eco MG1655 rph+ lac+ ΔmallI::frt gtrA::P58-mCherry:parB-SpR<br>codA::parBMt1-lacOsym-lacOCS-frt recA-recX::recAmwg-SYFP2-frt<br>DE(recB)::frt / p15a-wtCas9deg-SOS-amp + psgRNA_CS1 |
| EL2837 | Eco MG1655 rph+ recA::ALFA-recAmwg::SYFP2-frt-kan-frt<br>codA::parBMt1-lacOsym-lacOCS-frt / p15a_SceIdeg_amp |
| EL2858 | Eco MG1655 rph+ lac+ ΔmallI::frt gtrA::P58-mCherry:parB-SpR<br>codA::parBMt1-lacOsym-lacOCS-frt recA::syfp2-frt |
| EL2860 | Eco MG1655 rph+ lac+ ΔmallI::frt gtrA::P58-mCherry:parB-SpR<br>codA::parBMt1-lacOsym-lacOCS-frt recA-recX::recAmwg-SYFP2-frt /<br>p15a-dCas9deg-ampR / psgRNA_CS1 |
| EL2861 | Eco MG1655 rph+ lac+ ΔmallI::frt gtrA::P58-mCherry:parB-SpR<br>codA::parBMt1-lacOsym-lacOCS-frt recA:ALFA-frt/ p15a-dCas9deg-<br>ampR / psgRNA_CS1 |
| JW2788<sup>36</sup> | ΔrecB::KmR |
| JW3627<sup>36</sup> | ΔrecG::KmR |
| JW1852<sup>36</sup> | ΔrucC::KmR |
| EL1605<sup>37</sup> | Eco BW25993 DELphi80::yciI INSlacY::ypet INSintC::tetR-dcas9-spR<br>INSaraBAD::T7rnep-tetA DELaraB |
| EL1743 | MG1655 dnaC2 ΔmdoB::aph frt |

**Table S2.**
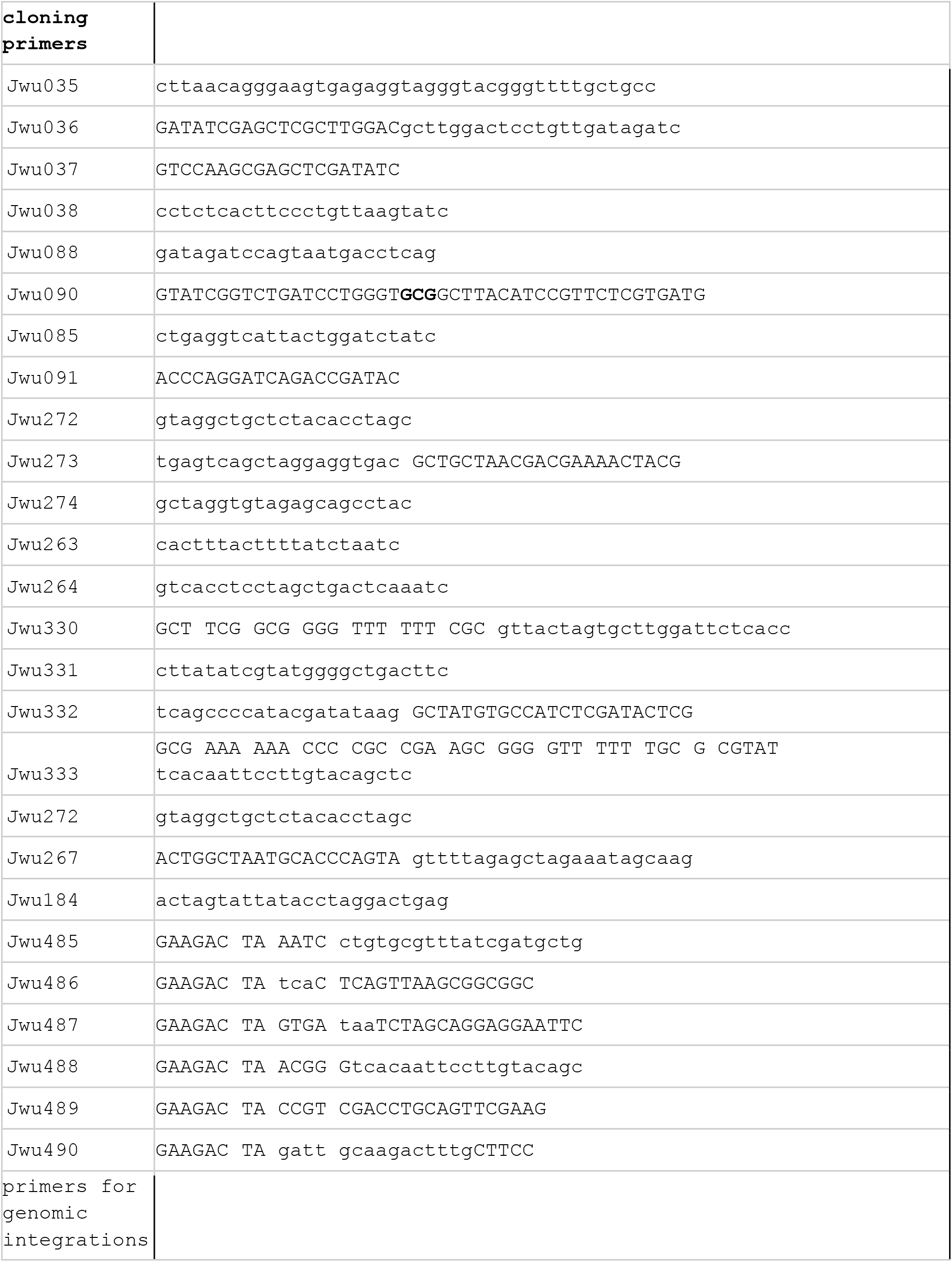
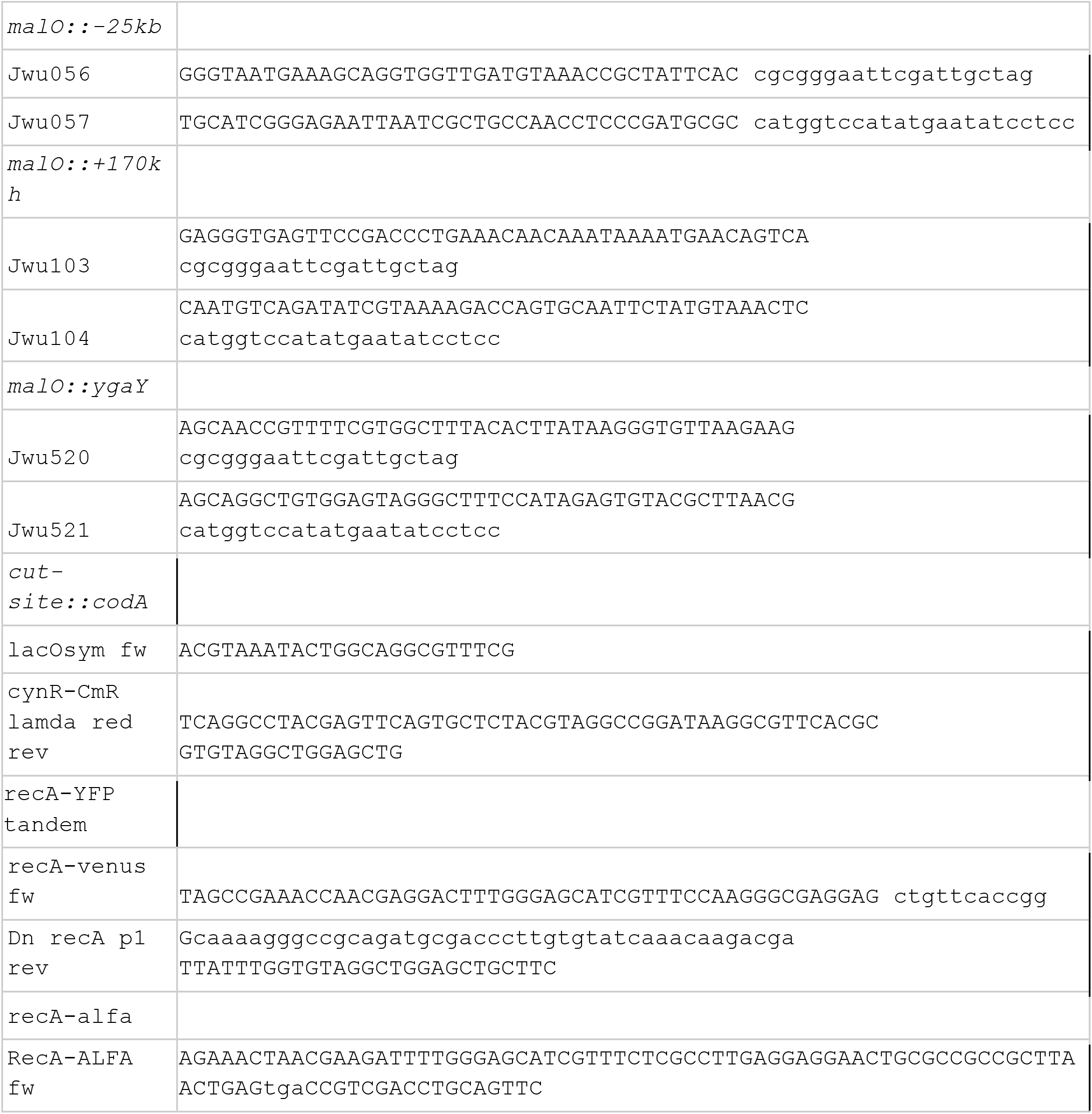
Primers used in this study

